# Sphingosine-1-Phosphate (S1P) analogue Phyto-sphingosine-1-Phosphate (P1P) improves the *in vitro* maturation efficiency of porcine oocytes via regulation of oxidative stress and apoptosis

**DOI:** 10.1101/499798

**Authors:** Kyu-Mi Park, Jae Woong Wang, Yeong-Min Yoo, Ji Eun Jang, Myeong Jun Choi, Sang Hwan Hyun, Kyu Chan Hwang, Eui-Bae Jeung, Yeon Woo Jeong, Woo Suk Hwang

## Abstract

Phytosphingosine-1-Phosphate (P1P) is a signaling sphingolipid regulating various physiological activities. Yet, little is known of the effect of P1P in the context of reproduction. As such, we aimed to investigate the influence of P1P on oocyte maturation during porcine *in vitro* maturation (IVM). Here we report the expression of S1PR1-3 among P1P receptors (S1PR1-4) in cumulus cells and oocytes. When P1P was treated by concentrations 10 nM, 50 nM, 100 nM, and 1000 nM during IVM, Metaphase II rate was significantly increased in 1000 nM (=1 μM) P1P treatment group. Maturation rate improvement by P1P supplementation was only observed in the presence of EGF. Oocytes under the influence of P1P decreased intracellular ROS levels yet did not show significant differences in GSH levels. In our molecular studies, P1P treatment up-regulated gene expressions involved in cumulus expansion (*Has2* and *EGF*), antioxidant enzyme (*SOD3* and *Cat*), and developmental competence (*Oct4*) while activating ERK1/2 and Akt signaling. P1P treatment also influenced oocyte survival by shifting the ratio of *Bcl-2* to *Bax*, while inactivating JNK signaling. We further demonstrated that oocytes matured with P1P significantly displayed not only higher developmental competence (cleavage and blastocyst formation rate), but also greater blastocyst quality (total cell number and the ratio of apoptotic cells) when activated via parthenogenetic activation (PA) and *in vitro* fertilization (IVF). Despite low levels of endogenous P1P found in animals, exogenous P1P was able to influence animal reproduction as shown by increased porcine oocyte maturation as well as preimplantation embryo development.

## Introduction

The current state of knowledge on embryonic development stems in part from numerous past studies using oocytes of industrial animals as a model system. Pigs have served as valuable animal resources for advancing biomedical research as pigs share many similarities to humans in terms of anatomy, physiology, and genetics [1, 2]. Immature oocytes obtained from these ovaries are often subject to *in vitro* production (IVP), which is a process composed of *in vitro* maturation (IVM) to derive mature oocytes and subsequently *in vitro* culture (IVC) to derive blastocysts. In comparison to other species, porcine IVM period is lengthy, requiring approximately 40 hours for maturation. As a consequence of a lengthy culture period, a notable degree of apoptosis and excessive reactive oxygen species (ROS)-induced oxidative stress (OS) accumulation may be observed. This added vulnerability has helped establish porcine IVP as a suitable model for studying apoptosis and ROSin reproduction. Numerous past studies have aimed to minimize this phenomenon, experimenting with various supplements [3–6].

Amongst various biologically active compounds that may potentially influence IVP, we concentrated our attention on a bioactive lipid mediator called Phyto-sphingosine-1-Phosphate (P1P; PhS1P), a phosphorylated form of phytosphingosine abundantly observed in plants and less in animals [7, 8]. Recently, P1P has seen a rise in popularity as a cosmetic substance owing to technological advancements allowing mass production, lowered costs, and increased solubility [9, 10]. P1P shares many similarities with its analog Sphingosine-1-Phosphate (S1P), both structurally and functionally. Two natural S1P analogues exist - P1P (PhS1P; Phyto-sphingosine-1-Phosphate) and DhS1P (Dihydrosphingosine-1-Phosphate). P1P has a similar structure to DH-S1P with an additional hydroxyl group at C-4 present at the sphingoid long-chain base being the exception (Fig 1). Likely owing to this structural similarity, P1P has been previously shown to interact with S1P receptors, a family of G-protein coupled receptors. Specifically, P1P binds to all S1P receptors with particularly high affinity to S1PR1 and S1PR4 while S1P binds to all five S1PR1-5 [11, 12].

**Fig 1.**
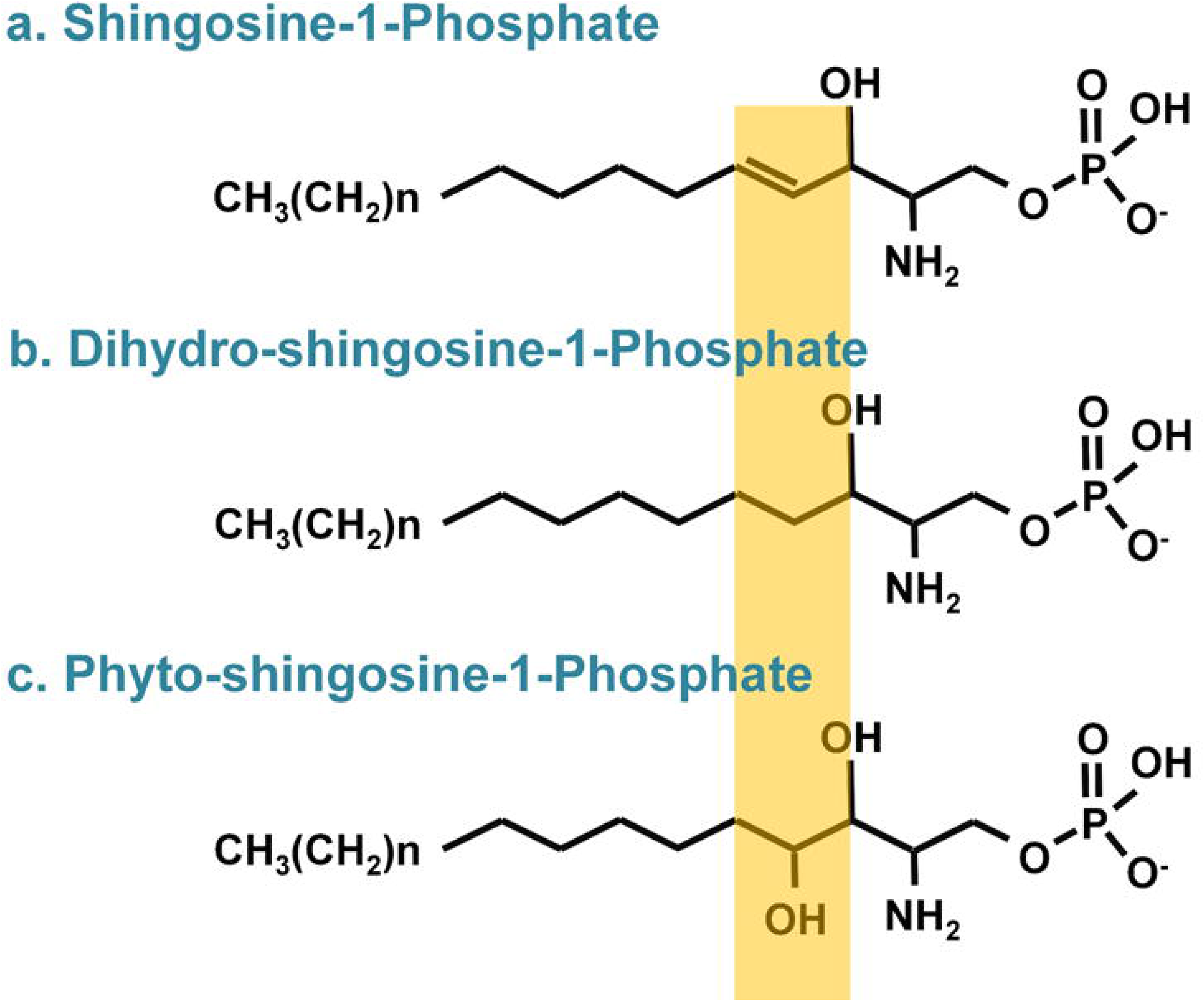
Structures of Sphingosine-1-Phosphate (S1P), Dihydrosphingosine-1-phosphate (DhS1P) and Phyto-sphingosine-1-Phosphate (P1P)

In general, S1P signaling has been demonstrated to activate phospholipase C (PLC), phospholipase D (PLD), mitogen-activated protein kinase (MAPK)/extracellular signal-regulated kinase (ERK), and protein kinase C (PKC) pathways [13–18]. Through combinatorial activation of such pathways, S1P carries out different functions in various biological contexts [19–21]. Like S1P, P1P also has a similar effect on cell proliferation, migration, and inhibition of OS-induced damage. P1P attenuates OS-induced cell growth arrest by regulating Akt and Jun amino-terminal kinases (JNK) [22], stimulates chemotactic migration through modulating p38 kinase and PI3K [23], and promotes chondrocyte proliferation via ERK-mediated pathway [24].

To postulate potential actions of P1P, studies have referred to previous results from P1P’s close analogue S1P. In reproductive systems, the roles of S1P have been illustrated in many studies. In human, the concentration of S1P in follicular fluid (FF) is positively correlated with pregnancy outcome [25], and in mouse and rat, S1P protected ovaries from damages acquired from external stimuli like chemotherapy and radiation therapy [26–28]. In the absence of S1P receptors, partial embryonic lethality and vascular abnormalities were found in mice, indicating the role of S1P receptors on embryonic angiogenesis [29]. As a model for testing S1P’s actions on oocyte maturation in vitro, several studies have carried out IVM supplemented by varying concentrations of S1P. In murine and porcine studies, S1P treatment increased developmental competence as observed by improved blastulation rate and/or decreased blastocyst apoptosis [30, 31]. In bovine, S1P improved survival and competence of oocytes exposed to heat shock during IVM [32].

Molecular studies have shown anti-apoptotic effects of S1P are exerted through inhibition of OS-induced apoptosis in granulosa cells by phosphorylation of the PI3K/Akt pathway [33]. There are numerous other substances that activate the PI3K/Akt pathway, which is commonly implicated in cell proliferation, differentiation, and anti-apoptosis. One of such factors, epidermal growth factor (EGF) has been shown to have a close relationship to P1P/S1P signaling in non-reproductive systems. In fibroblast cells, P1P displays synergistic relationship with EGF via p-Akt and p-ERK activation, promoting wound healing and cell proliferation [34, 35]. In vascular smooth muscles, S1P stimulates epidermal growth factor receptor (EGFR) expression [36] whereas in gastric cancer cells, S1P induces the release of EGF, an activator of EGFR signaling [37]. Considering EGF is a well-studied factor in reproduction shown to improve meiotic resumption, verification of the synergistic relationship between EGF and P1P in reproduction is necessary. Despite the advancements made in the knowledge of S1P-mediated signaling, there is a lack of studies examining the effects of its close analog P1P, which is an evolutionarily-conserved, yet to be highlighted binding partner of S1P receptors.

In this study, we aimed to study the effects of P1P in the context of reproduction by P1P supplementation during IVM. Upon verification of the presence of S1P receptors expressions in COCs, we explored the dose-dependent effects of P1P on nuclear and cytoplasmic maturation. The synergistic effect of EGF with P1P during IVM was also examined. To support our observation, protein activations and gene expressions implicated in cumulus cells expansion, enzymatic antioxidants, developmental competence, and apoptosis were assessed. Thereafter, the effects of P1P treatment during IVM on subsequent pre-implantation development is shown by cleavage rate, blastocyst rates, and apoptotic patterns. Together, this study highlights the roles of P1P supplementation during IVM on various parameters of meiotic resumption and developmental competence.

## Materials and methods

P1P was obtained from Phytos Co. (Korea). Unless specified otherwise, all chemicals and reagents were purchased from Sigma-Aldrich Chemical Company (USA).

### IVM

Ovaries collected from a local abattoir were transported in saline solution at temperatures 32-35 °C. Cumulus-oocyte complexes (COCs) from antral follicles 3-6mm were aspirated using a 10 ml disposable syringe. Aspirated COCs were washed in HEPES-buffered Tyrode’s medium (TLH) with 0.05 % (w/v) polyvinyl alcohol (TLH-PVA). Only compact COCs and homogeneous cytoplasm were selected for the experiment. The base medium used for all subsequent experiments consisted of TCM199 (Invitrogen Corporation, USA) supplemented with 1 μg/ml insulin, 10 ng/ml EGF, 75 μg/ml kanamycin, 0.6 mM cysteine, 0.1 % (w/v) polyvinyl alcohol (PVA), 0.91 mM sodium pyruvate. 60 COCs per experimental group were matured in 500 μl of IVM medium for a total of 42 hours at 39 °C for 20 h under 5 % CO_2_ following a two-step culture system. During the first 22 hours of culture, 10 IU/ml hCG and 10 IU/ml eCG were supplemented to the base medium, and the remaining 20 hours of culture did not contain hCG nor eCG. During all culture period, experimental groups were treated with various concentration of P1P : 0 (control), 10, 50, 100, 1000 nM (1 μM).

Furthermore, to demonstrate the correlation between EGF and P1P, additional IVM experiment was performed in the absence or presence of 10 ng/ml EGF, depending on the experimental design.

### Reverse transcription polymerase chain reaction (RT-PCR)

To analyze P1P receptors expression, messenger RNA (mRNA) and genomic DNA (gDNA) were extracted from 150 post-IVM oocytes and denuded cumulus cells using Dynabeads kit (Invitrogen Corporation, USA) and G-DEX kit (Intron Biotechnology, Republic of Korea) following the manufacturer’s instructions. Complementary DNA (cDNA) synthesis was carried out using LaboPass^TM^ synthesis kit (Cosmo Genetech, Republic of Korea). The RT-PCR reactions consisted of incubation for 5 min at 95 °C (initial denaturation), followed by 40 cycles of 30 sec at 95 °C (denaturation), 30 sec at 58 °C (annealing), 30 sec at 72 °C (extension). The PCR products were electrophoresed on a 1.5 % agarose gel followed by visualization under UV light. The primer sequences used for the detection of P1P receptors are presented in Table 1.

**Table 1.**
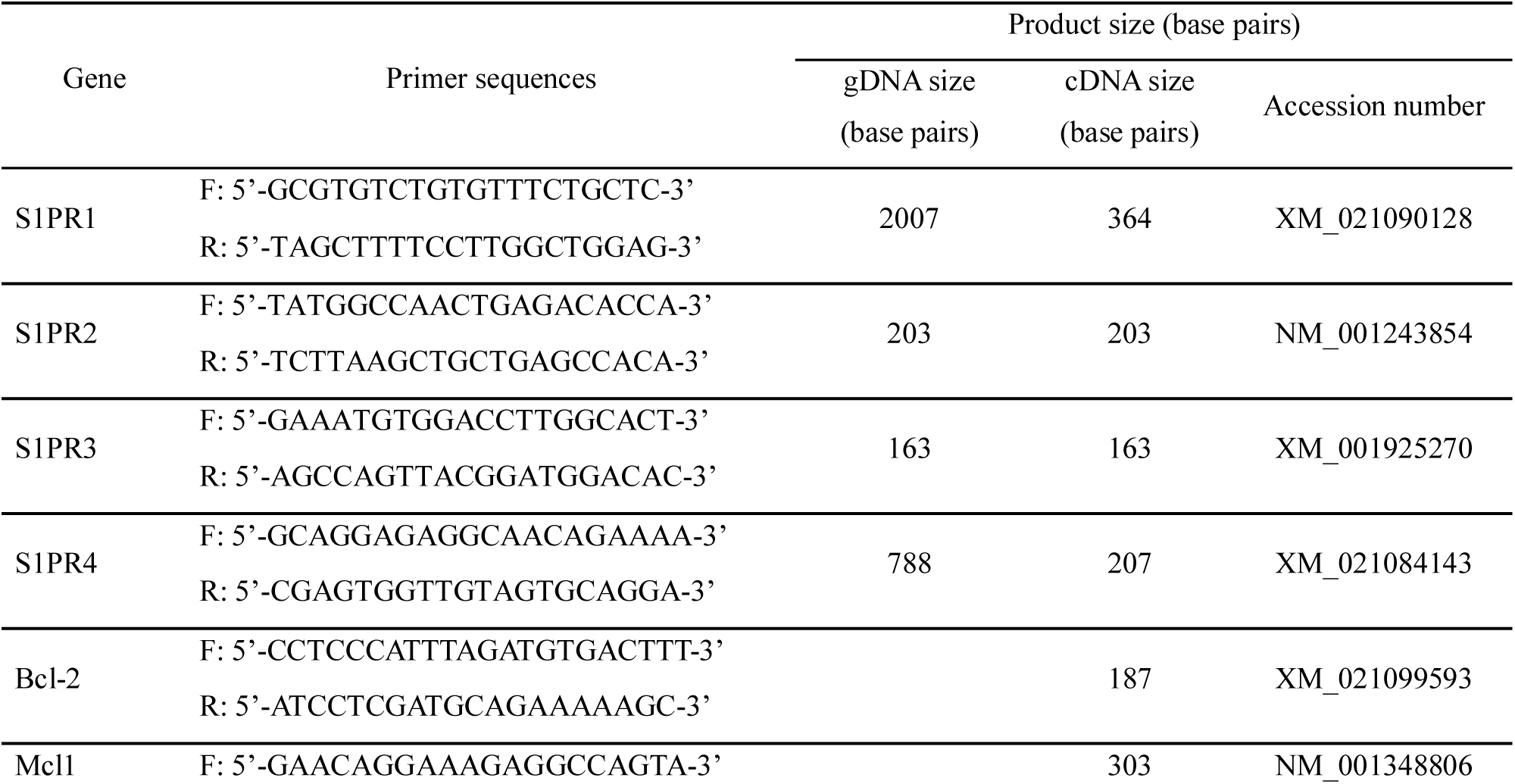
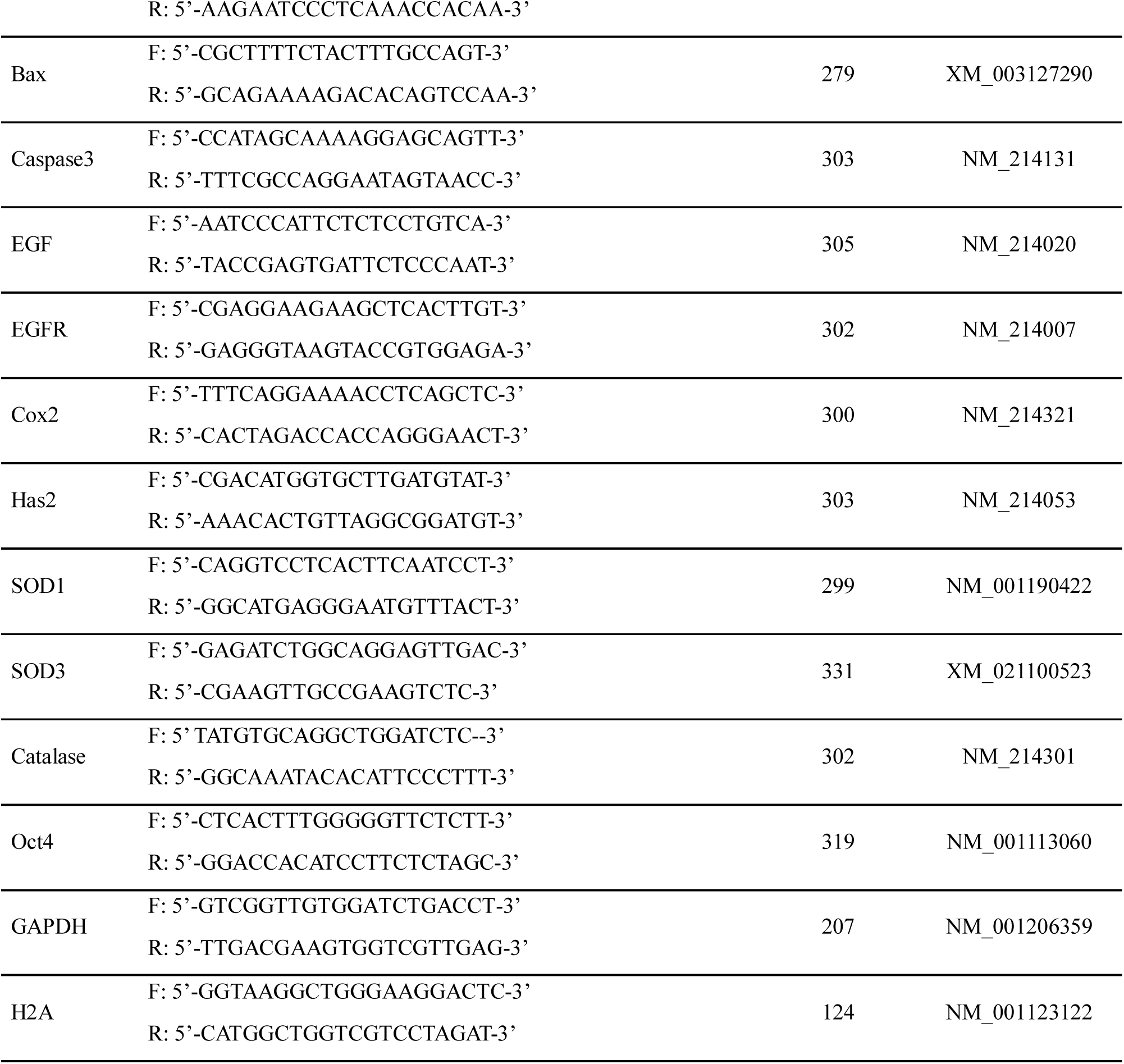
Primer Sequences for analysis of mRNA gene expression

### Assessment of chromatin configuration

To assess nuclear maturation, post-IVM COCs from each group were denuded by repeated pipetting in TLH medium supplemented with 0.1 % hyaluronidase. The denuded oocytes were fixed in 4 % paraformaldehyde and stained for 10 min with 10 μg/mL Hoechst 33342 diluted in TLH-PVA. The stained oocytes were examined under a fluorescence microscope (TE300, Nikon Corp., Japan) and classified as previously described [38]: germinal vesicle, germinal vesicle breakdown, metaphase I, anaphase-telophase I, or metaphase II. The experiment was replicated four times (n=4).

### Measurement of intracellular reactive oxygen species (ROS) and glutathione (GSH) levels

Intracellular ROS and GSH levels of MII stage oocytes post-IVM were measured as previously described [39]. Briefly, post-IVM MII stage oocytes from each group were incubated for 30 min in either 10 μM 2’,7’-dichlorodihydrofluorescein diacetate (H2DCFDA; Invitrogen Corporation) or 10 μM 4-chloromethyl-6.8-difluoro-7-hydroxycoumarin (Cell Tracker Blue; Invitrogen Corporation) to identify intracellular GSH levels and ROS levels, respectively. After incubation, oocytes were rinsed in Dulbecco’s phosphate buffer saline (DPBS) containing 0.1 % (w/v) PVA (PVA-PBS) and were observed under an epifluorescence microscope (TE300; Nikon, Japan) equipped with 370 nm and 460 nm excitation filters. The fluorescence intensity was analyzed by Adobe Photoshop software (Version 10.0, USA). The experiment was replicated three times (n=3).

### Quantitative real-time polymerase chain reaction (Real-time PCR)

Gene expression levels related to cumulus expansion, enzymatic antioxidants, developmental competence, and apoptosis were quantified by real-time PCR. The method used for cDNA synthesis from mRNA extracted from cumulus cells and oocytes was identical to that used for detection of P1P receptors expression described previously above. All polymerase chain reactions were performed in 20 μL reaction volume containing 1 μL cDNA template, a specific primer, and SYBR Green Q master mix (Cosmo Genetech). The program for PCR amplification consists of inactivation (10 min at 95 °C), followed by 40 cycles of denaturation (30 sec at 95 °C), annealing (30 sec at 62 °C) and extension (30 sec at 72 °C). The primer sequences used are summarized in Table 1. The expression level of each gene was normalized to *H2A* expression level by relative threshold cycle (CT). The relative expression (R) was calculated by the equation, R = 2^-[Δ Ct sample – Δ Ct control]^. The experiments were repeated four and three times each for the analyses of cumulus cells and oocytes, respectively.

### Western blotting

Western blot analyses were performed as previously described unless stated otherwise [40]. 1 × 10^5^ cumulus cells from the control and 1 μM P1P treatment group after IVM were sampled and lysed using EzRIPA Lysis kit (WSE-7420; ATTO Corporation, Republic of Korea) following the manufacturer’s instructions. Protein concentrations were validated using the BCA (Bicinchoninic acid) ^TM^ Protein Assay kit (23225; Thermo Fisher Scientific), and each protein sample (10 μg) was separated on 12 % SDS-PAGE gels and transferred onto polyvinylidene fluoride (PVDF) membranes. The membranes were incubated at 4°C with primary antibodies against ERK1/2 and p-ERK1/2 (9101s, Cell Signaling Technology, USA, 1:1000); Akt and p-Akt (9271s, Cell Signaling Technology, 1:1000); SAPK/JNK and p-SAPK/JNK (9251s, Cell Signaling Technology, 1:1000) for overnight. After overnight incubation, the membranes were incubated with anti-rabbit IgG-conjugated horseradish peroxidase secondary antibody (Cell Signaling Technology) for 2 hours at room temperature. Immunoreactive proteins were visualized using X-ray film and the bands were imaged using a scanner. ImageJ software (version 1.37; Wayne Rasband, NIH, USA) was used for the measurement of optical density and the data were normalized to internal control. All experiments were repeated eight times (n=8).

### Parthenogenetic activation (PA) and IVC

For parthenogenetic activation (PA), denuded mature oocytes were washed twice and activated by an electrical pulse of 120 V/mm for 60ms in 260 mM mannitol solution supplemented with 0.1 mM CaCl_2_ and 0.05 mM MgCl_2_ by an electrical pulsing machine (LF101; Nepa gene, Japan). Activated oocytes were incubated in IVC media (porcine zygote medium 3; PZM3) containing 5 μg/mL of cytochalasin B for 4 h. After the incubation period, post-activation oocytes were washed twice and cultured in 30 μL droplets of PZM3 (10 embryos/droplet) for 7 days under 90 % N_2_, 5 % O_2_ and 5 % CO_2_ at 39°C. The cleavage (PA-CL) rates and patterns of all groups were recorded on day 2 after PA. The blastocyst (PA-BL) formation rates and patterns were, recorded on day 7 after PA. The experiment was repeated four times (n=4).

### *In vitro* fertilization (IVF)

After IVM, cumulus cells were removed by mechanically pipetting with 0.1% hyaluronidase and oocytes with the first polar body extruded were chosen for IVF. After washing with modified Tris-buffered medium (mTBM), groups of 15 oocytes were placed in 40 μL droplets of mTBM covered with mineral oil. Semen sample supplied by Darby Genetics Inc. (Department of Livestock Research, Republic of Korea) previously stored at 17 °C were rinsed by centrifugation at 2000 × g for 3 min in DPBS supplemented with 0.1 % BSA. The 15 oocytes were incubated with 5 × 10^5^ sperm/mL spermatozoa in 45 μL droplets of mTBM under 5 % CO_2_ at 39 °C. After 20 min, loosely bound sperms were detached by gentle pipetting. Next, the gametes were then rinsed in mTBM and cultured to a droplet of mTBM without sperm for 5 h under 5 % CO_2_ at 39 °C. After 5 h, the gametes were washed twice, cultured, and evaluated the cleavage (IVF-CL) rates and blastocyst (IVF-BL) rates as in the IVC method described above. The experiments were repeated four times (n=4).

### TUNEL assay and total cell count

All blastocysts after PA and IVF washed in PVA-PBS and subsequently fixed in DPBS containing 4 % paraformaldehyde at 4 °C for 1 h. Next, Fixed blastocysts rinsed in PVA-PBS and permeabilized overnight in DPBS containing 0.1 % (w/v) Triton X-100 at 4 °C. Fixed blastocysts were washed in PVA-PBS and incubated in droplet mixed 9 μl fluorescein-conjugated dUTP and 1 μl terminal deoxynucleotidyl transferase (TUNEL, In Situ Cell Death Detection Kit, Roche, Germany) at 38 °C for 1 h in the dark. After series of washings, all blastocysts were counterstained in DPBS supplemented with 10 μg/mL Hoechst 33342 for 5 min, washed, placed on a slide glass, compressed with a coverslip, and observed under fluorescence microscope.

### Statistical analysis

Statistical analyses were performed by SPSS (Version 17.0, USA). Using one-way ANOVA, followed by Duncan’s multiple range test, the percentage data of maturation, cleavage and blastocyst rates, the total cell numbers of the blastocyst and the ratio of apoptotic cells were compared. Relative gene expression levels and protein expression levels were analyzed by the student *t*-test. Values are presented as the mean ± SEM. Statistically, significant differences are indicated by appropriate symbols or letters described in figure legends.

## Results

### S1P Receptors are expressed in cumulus cells and oocytes

Previous studies have shown that P1P acts by binding to S1P receptors present on cell surfaces [11, 12]. Prior to investigating the effects of P1P on porcine meiotic resumption, gene expression of S1P receptors *S1PR1-4* in cumulus cells and oocytes were identified. The presence of all four *S1PR1-4* genes in the porcine genome was confirmed by PCR (Fig 2A). To further test for the presence of *S1PR1-4* RNA transcript, RT-PCR was conducted on cumulus cells and oocytes. Among P1P receptors, *S1PR2* and *S1PR3* transcripts were expressed in both cumulus cells and oocytes while *S1PR1* was expressed only in cumulus cells and not in oocytes. S1PR4 expression was detected in neither cumulus cells nor oocytes (Figs 2B and C). As such, the expression of P1P receptors S1PR1-3 was confirmed, leading to the hypothesis that P1P may affect in vitro maturation of porcine oocytes via S1PRs.

**Fig 2.**
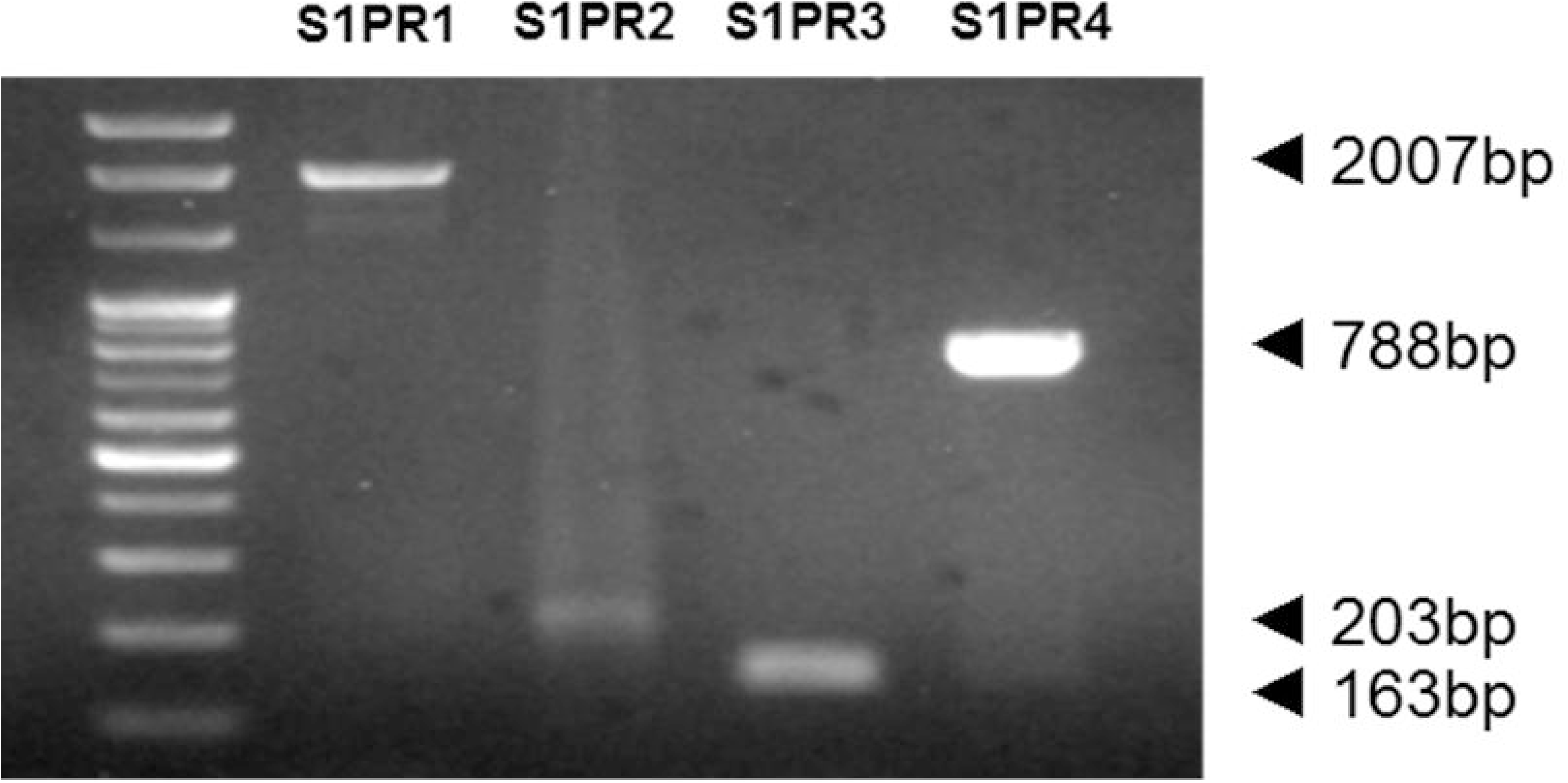
Identification of P1P receptors (S1PR1-R4) gene expression in matured cumulus cells and oocytes. (A) Genomic DNA (gDNA) and (B, C) complementary DNA (cDNA) polymerase chain reaction analysis in cumulus cells and oocytes

### P1P enhances nuclear maturation during porcine in vitro maturation

To test for the potential effects of P1P on porcine oocyte meiotic maturation, different concentrations of P1P were supplemented during IVM, and oocytes were assessed for nuclear stage post-IVM. The rate of MII stage oocytes in the 1000 nM (=1 μM) group was significantly higher than the MII rates of the control, 10 nM, and 50 nM group (p<0.05). In contrast, no significant differences were found in MII rates between the 100 nM and 1000 nM P1P groups (Table 2). To investigate whether higher concentrations would have further beneficial effects on meiotic maturation, the effects of 5 μM P1P was additionally tested. Unlike at 1 μM P1P, supplementation of 5 μM P1P did not significantly improve meiotic maturation compared to non-treated control (S1 Table). In short, at the concentrations tested, 100 nM to 1000 nM P1P supplementation had the most beneficial effects in promoting meiotic maturation.

**Table 2.**
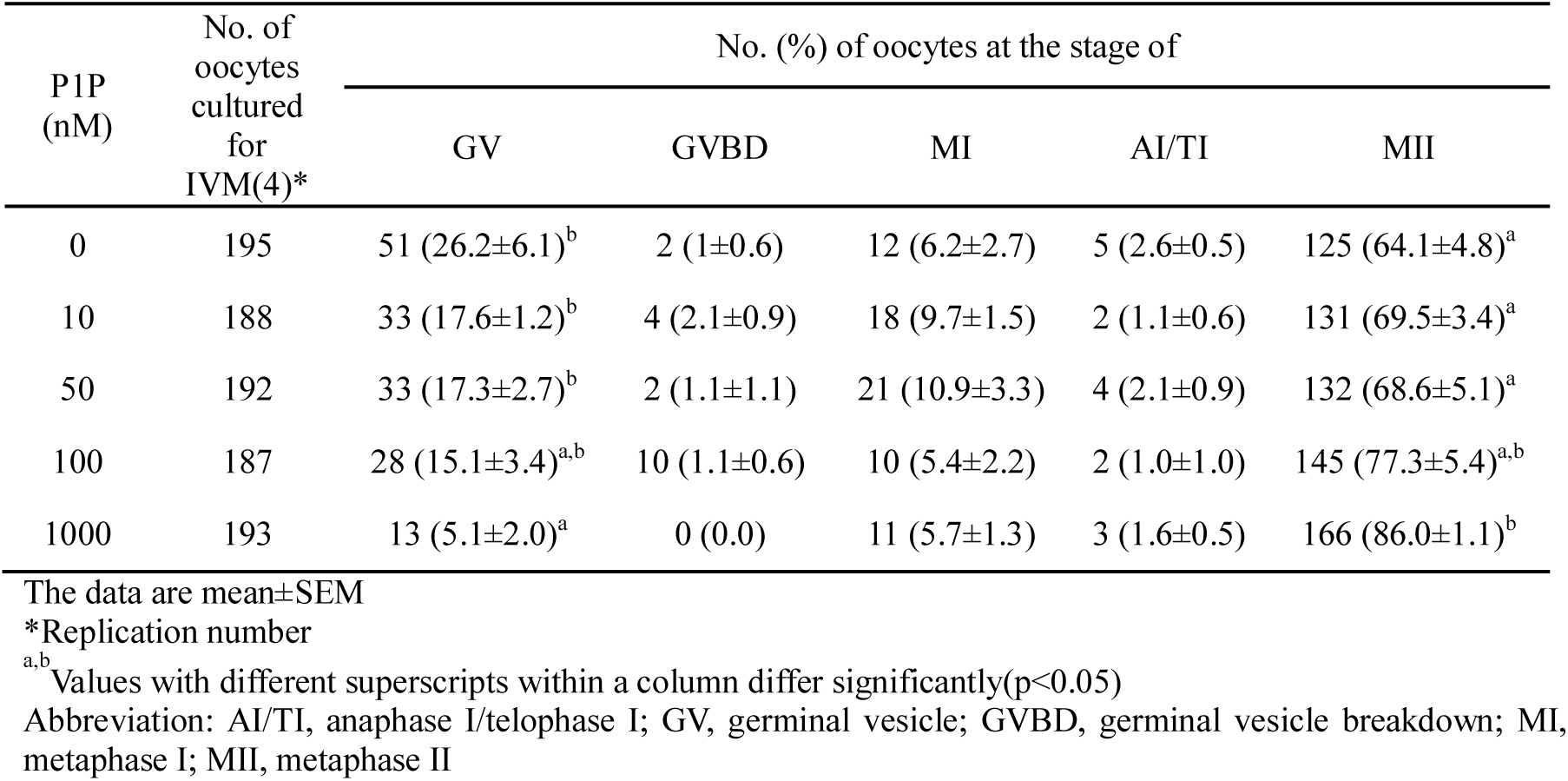
Effects of P1P treatments on nuclear maturation in porcine oocytes

### P1P depends on a synergistic relationship with EGF for promoting meiotic maturation in vitro

The effects of P1P on meiotic maturation of porcine oocytes has prompted us to further characterize the molecular pathways responsible for the effects observed above. Previous studies have reported synergistic involvements of growth factors such as EGF and P1P in molecular pathways important for fibroblast cells [34, 35]. Considering the widespread use of EGF in IVM, we investigated whether the effects of P1P in meiotic maturation may be in part due to synergistic effects with EGF by experimenting the effects of P1P in the presence or absence of EGF. In the presence of EGF, MII rate was significantly higher when supplemented with 1 μM P1P as shown previously demonstrated. In contrast, P1P supplementation did not significantly affect meiotic maturation in the absence of EGF (Table 3). Taken together, these observations suggest that the effects of P1P on promoting meiotic maturation depends in part on the synergistic relationship with EGF.

**Table 3.** Effects of P1P treatments during IVM according to absence EGF on nuclear maturation

| PIP<br>(μM) | | No. of<br>oocytes<br>cultured<br>for<br>IVM(4)* | No. (%) of oocytes at the stage of | | | | |
| --- | --- | --- | --- | --- | --- | --- | --- |
|  |  |  | GV | GVBD | MI | AI/TI | MII |
| EGF(-) | 0 | 267 | 41 (15.4±1.9) | 2 (0.8±0.4) | 35 (12.9±1.6)<sup>c</sup> | 6 (2.2±0.8) | 183 (68.7±2.0)<sup>a</sup> |
| | 1 | 268 | 31 (11.3±3.9) | 1 (0.3±0.3) | 29 (11.1±2.1)<sup>b,c</sup> | 3 (1.1±0.7) | 204 (76.2±2.5)<sup>a,b</sup> |
| EGF(+) | 0 | 265 | 35 (12.8±7.6) | 0 (0.0) | 21 (8.0±1.0)<sup>a,b</sup> | 4 (1.5±0.7) | 205 (77.6±3.3)<sup>b</sup> |
| | 1 | 268 | 18 (6.6±1.7) | 0 (0.0) | 16 (5.8±0.9)<sup>a</sup> | 1 (0.4±0.4) | 233 (87.2±2.5)<sup>c</sup> |

The data are mean ± SEM
\*Replication number
<sup>a,b</sup> Values with different superscripts within a column differ significantly (p<0.05)
Abbreviation: AI/TI, anaphase I/telophase I; GV, germinal vesicle; GVBD, germinal vesicle breakdown; MI, metaphase I; MII, metaphase II

### P1P supplementation reduces intracellular ROS level independent of GSH

Besides nuclear maturation, another important factor to consider when assessing oocyte maturation is cytoplasmic competence. Intracellular ROS and GSH levels are commonly studied parameters for assessing oocyte cytoplasmic quality [4]. We characterized the intracellular GSH and ROS levels of MII oocyte groups treated with various concentrations of P1P during IVM. In Fig 3, the GSH levels of oocytes show no significant variation between control and treatment groups (p<0.05). In comparison, the ROS levels were reduced in a dose-dependent manner. The intracellular ROS levels of oocytes in 100 nM and 1000 nM P1P treatment groups were decreased significantly compared to that of the control group (p<0.05). These findings suggest that P1P supplementation decreases ROS levels in porcine oocytes during IVM by mechanisms possibly independent of GSH levels.

**Fig 3.**
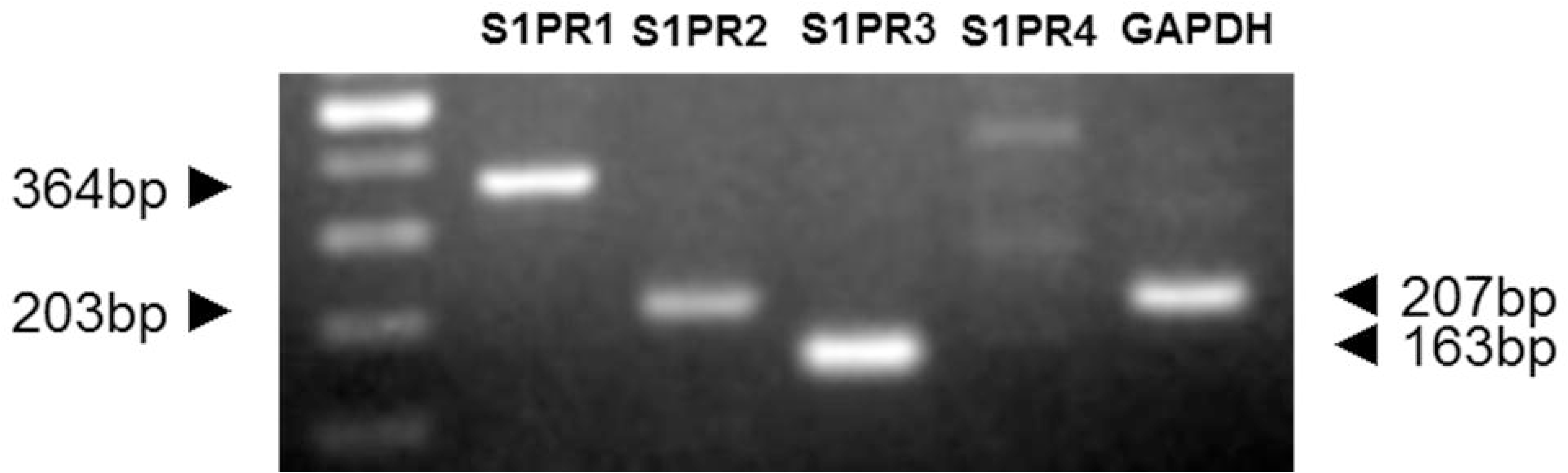
Epifluorescent photomicrographic images of in vitro matured porcine oocytes. (A) Oocytes were stained with Cell Tracker Blue (a–d) and 2’, 7’-dichlorodihydrofluorescein diacetate (H2DCFDA) (e–h) to detect the intracellular levels of glutathione (GSH) and reactive oxygen species (ROS). Metaphase II (MII) oocytes treated 0nM (a, f), 10nM (b, g), 50nM (c, h), 100nM (d, i) and 1000nM P1P (e, j). (B) Effect of P1P treatment on the intracellular GSH and ROS levels in porcine oocytes matured in vitro. GSH samples, N=24; ROS samples, N=24. This experiment was replicated three times. a, b, c Values with different superscripts within a column differ significantly (p<0.05).

### P1P affects the mRNA transcript levels associated with cumulus expansion, enzymatic antioxidants, and apoptosis

To further investigate the molecular pathways affected by P1P during porcine IVM, we analyzed the expression of genes important for meiotic resumption separately in cumulus cells and oocytes. In post-IVM cumulus cells, the transcript levels of expansion-related genes *EGF* and *HAS2* in 1 μM (=1000 nM) P1P treatment group was significantly higher than in the control group (* p<0.05) while expressions of *EGFR* and *Cox2* were not significantly different. Upon examination of apoptosis-related genes, we found *Bcl-2* and *BAX* transcript levels in the treatment group were significantly increased (* p<0.05) while transcript level of *Mcl1* and *Cas3* were not significantly different (Fig 4A). In post-IVM denuded oocytes, treatment with P1P significantly increased the expression of enzymatic antioxidants *SOD3* and *Cat* (^†^ p=0.065) while the expression of *SOD1* was not significantly affected. The expression of developmental competence gene *Oct4* also was significantly increased in P1P treatment (^‡^ p=0.079). As in cumulus cells, a similar increase in *Bcl-2* expression level was found in oocytes (* p<0.05). In contrast, *Bax* expression was significantly decreased in oocytes (** p=0.0516), unlike in cumulus cells. P1P did not significantly affect *Cas3* and *Mcl1* expression compared to untreated control (Fig 4B).

**Fig 4.**
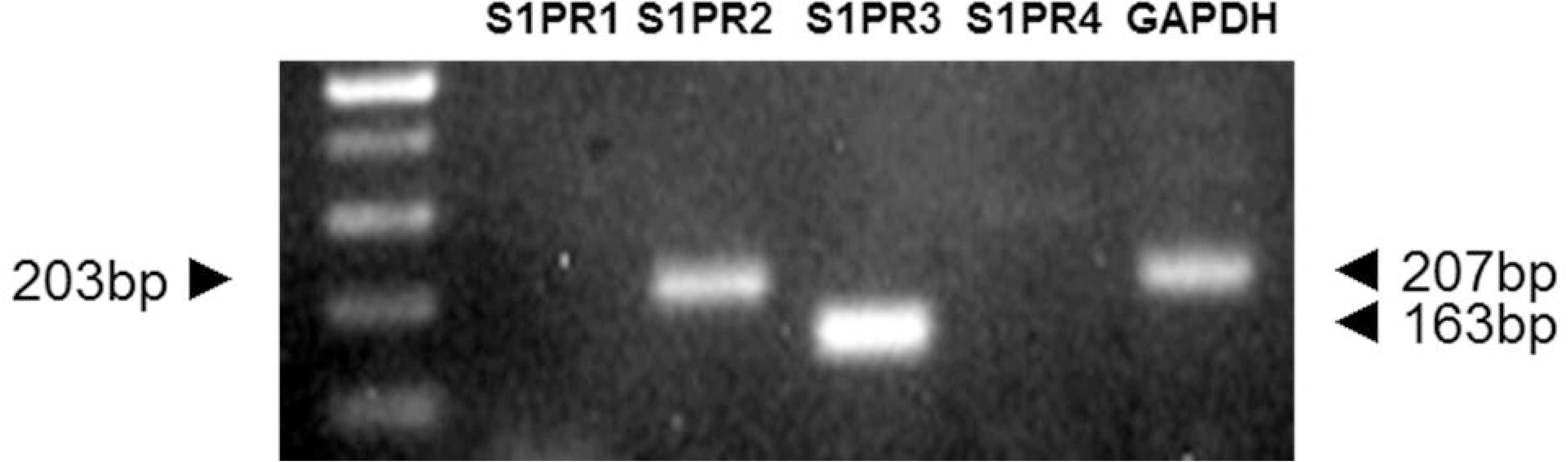
Real-time PCR analysis using cumulus cells and oocytes. We checked the mRNA transcripts in (A) cumulus cells and (B) oocytes supplemented with or without 1 μM P1P during IVM. (C) The ratios of BCL-2 to BAX in cumulus cells and oocytes. The expression of each product was quantified relative to that of the reference gene (H2A). CAS, Caspase 3; CAT, Catalase * p < 0.05; ** p = 0.0516; † p = 0.065; ‡ p = 0.079 as determined by t-test

To investigate the biological significance of the varying expression of apoptosis-related genes, we calculated the relative ratio of *Bcl-2* to *Bax*, which ratio has been previously illustrated to be a useful measure for predicting the balance between cell death and survival [41–43]. In our study, P1P supplementation led to significant increases in *Bcl-2* to *Bax* ratios in both cumulus cells and oocytes, favoring survival (* p<0.05) (Fig 4C). Together, P1P supplementation led to increases in expression of genes associated with cumulus expansion, enzymatic antioxidants, developmental competence, and survival.

### P1P induces phosphorylation of ERK1/2 and Akt but reduces phosphorylated JNK in cumulus cells

Previous studies demonstrated that P1P affects the activation of ERK1/2, Akt, and JNK in various cell type [22, 24]. In cumulus cells, the phosphorylation of ERK1/2, Akt, and JNK in cumulus cells has been shown to influence cumulus expansion and apoptosis during IVM [44, 45]. To investigate whether P1P improved meiotic maturation and oocyte quality through these pathways, western blot was performed to measure ERK1/2, Akt and SAPK/JNK protein expression and their phosphorylated form (Fig 5A). 1 μM P1P treatment led to significant increase in the expression of p-ERK1/2 and p-Akt compared to the control group whereas the expression of p-JNK was significantly decreased than that of control (p<0.05) (Fig 5B). These findings suggest that the presence of P1P during IVM led to the activation of ERK1/2 and Akt as well as the inactivation of JNK.

**Fig 5.**
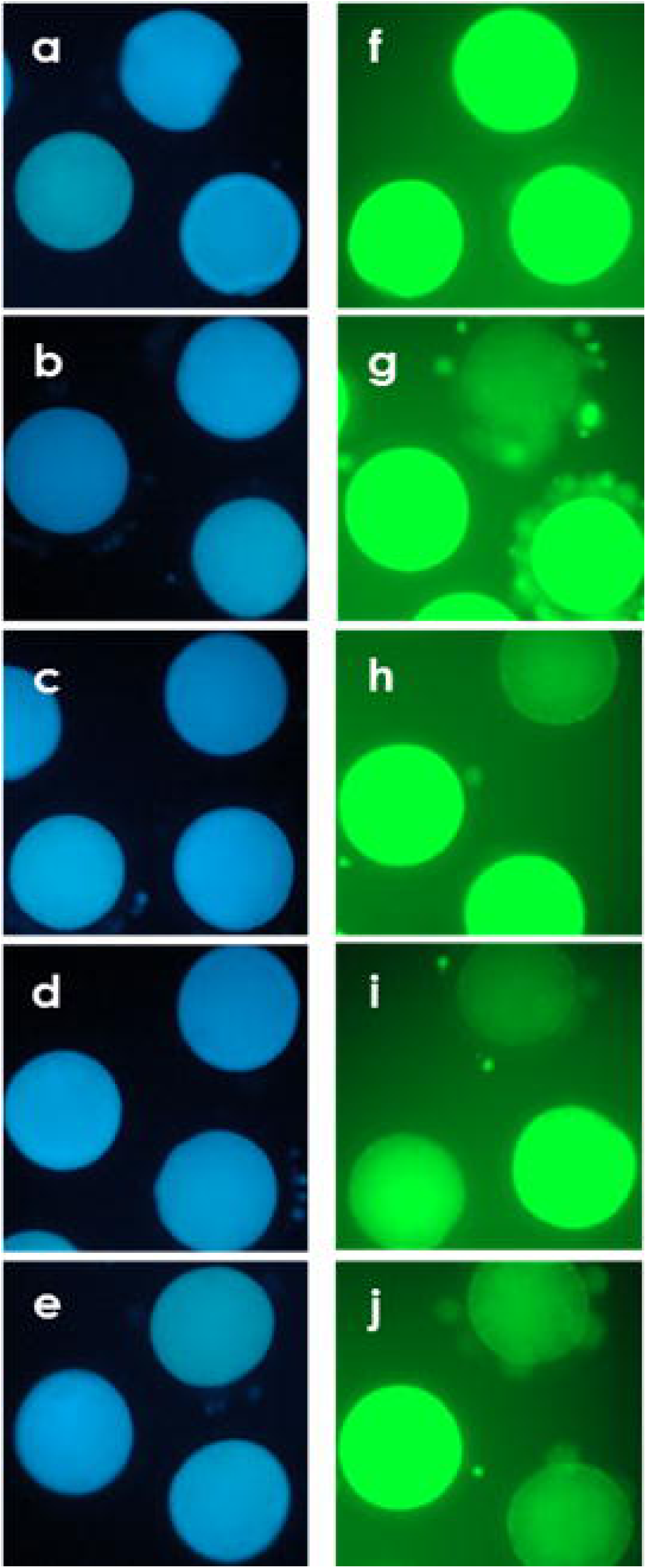
Phosphorylation of ERK 1/2, AKT, and JNK in the absence or presence of 1 μM P1P. (A) p-ERK 1/2, p-AKT and p-JNK expressions were analyzed by Western blot. (B) The relative amount of p-ERK 1/2, p-AKT and p-JNK were quantified as described in the method section. This experiment was replicated eight times. * p < 0.05 as determined by t-test

### P1P treatment during IVM period improved preimplantation embryo development and quality

To determine whether the improvement in meiotic resumption observed by P1P treatment is correlated to improvements in developmental competence, we assessed the subsequent embryonic development of PA and IVF embryos treated with different dosages of P1P during IVM. In the PA experiment, the PA-CL and PA-BL rate increased in the 100 nM and 1000 nM P1P group in comparison to 0, 10 and 50 nM groups (p<0.05) as shown in Table 4. However, no significant differences were observed in the PA-CL and PA-BL rates between 100 nM and 1000 nM group. In parallel, IVF experiment results presented significantly higher IVF-CL rates in the 1000 nM P1P group in comparison to 0, 10, 50 nM groups (p<0.05) (Table 5). Unlike IVF-CL rate, IVF-BL rate was significantly higher only in the 1000 nM group compared to groups treated with lower dosages (p<0.05).

**Table 4.** Effects of P1P treatments during *in vitro* maturation (IVM) on the embryonic development after parthenogenetic activation (PA)

| P1P<br>(nM) | No. of oocytes<br>cultured for IVM(4)* | No. (%) of embryos developed to |  |  |  |
| --- | --- | --- | --- | --- | --- |
| | | $\geq 2$-cell | | Blastocyst | |
| 0 | 177 | 118 | (66.8±1.4)<sup>a</sup> | 67 | (37.9±3.8)<sup>a</sup> |
| 10 | 170 | 118 | (69.7±3.6)<sup>a</sup> | 61 | (36.5±4.3)<sup>a</sup> |
| 50 | 181 | 126 | (69.5±1.3)<sup>a</sup> | 68 | (38.1±3.2)<sup>a</sup> |
| 100 | 185 | 139 | (74.8±2.3)<sup>a,b</sup> | 81 | (43.8±1.8)<sup>a,b</sup> |
| 1000 | 198 | 156 | (78.6±3.3)<sup>b</sup> | 101 | (51.0±2.2)<sup>b</sup> |

The data are mean ± SEM
\*Replication number
<sup>a,b</sup> Values with different superscripts within a column differ significantly(p<0.05)

**Table 5.** Effects of P1P treatments during *in vitro* maturation (IVM) on the embryonic development after *in vitro* fertilization (IVF)

| P1P<br>(nM) | No. of oocytes<br>cultured for IVM(4)* | No. (%) of embryos developed to |  |  |  |
| --- | --- | --- | --- | --- | --- |
| | | $\geq 2$-cell | | Blastocyst | |
| 0 | 173 | 110 | (62.9±3.2)<sup>a</sup> | 50 | (29.1±1.2)<sup>a</sup> |
| 10 | 173 | 114 | (65.8±2.1)<sup>a</sup> | 48 | (27.5±1.6)<sup>a</sup> |
| 50 | 180 | 111 | (62.1±2.1)<sup>a</sup> | 55 | (30.7±3.1)<sup>a</sup> |
| 100 | 208 | 147 | (70.7±5.0)<sup>a,b</sup> | 63 | (30.7±3.1)<sup>a</sup> |
| 1000 | 206 | 158 | (76.8±1.9)<sup>b</sup> | 85 | (40.4±3.2)<sup>b</sup> |

The data are mean ± SEM
\*Replication number
<sup>a,b</sup> Values with different superscripts within a column differ significantly(p<0.05)

The positive effects of P1P on increased embryo generation have prompted us to further characterize the quality and developmental potential of the blastocysts generated by quantifying the ratio of apoptotic cells to the total number of blastomeres (Fig 6A). In PA blastocysts, total blastocyst cell numbers significantly increased in the 1000 nM group in comparison to 0, 10 and 50 nM groups (p<0.05), whereas no significant differences were found between 100 nM and 1000 nM group (Fig 6B). The ratio of apoptotic cells in 1000 nM significantly decreased compared to that of 0 and 10 nM groups (p<0.05) whereas 50 nM, 100 nM, and 1000 nM groups did not show significant differences (Fig 6B). In parallel, IVF blastocysts in 1000 nM group presented significantly higher total blastocyst cell numbers and lower ratio of apoptotic cells in comparison to 0, 10, 50, and 100 nM groups (p<0.05) (Fig 6C). These findings possibly suggest that P1P treatment during IVM increases the efficiency of embryo generation and the quality of the generated embryos, especially in the 1 μM P1P treatment group.

**Fig 6.**
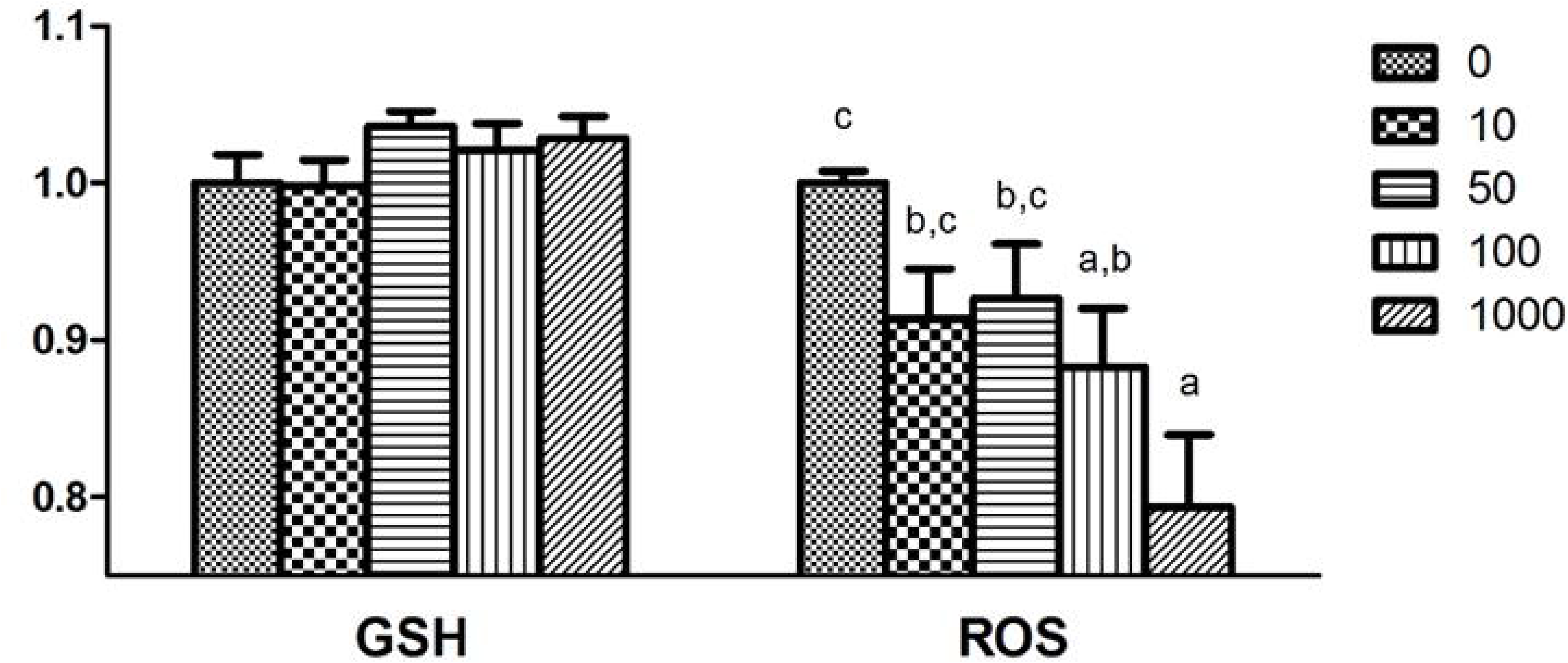
Hoechst and TUNEL staining of blastocysts. (A) Images of TUNEL and merge (TUNEL/HOCHEST 33342) stained blastocysts on day 7 after parthenogenesis (PA) and in-vitro fertilization (IVF). The scale bar is 50 μm. The number of total blastomeres and ration of apoptotic cells after (B) PA and (C) IVF. Within each bar with different letters (a,b and c) are significantly (P<0.05) different.

## Discussion

In the present study, we demonstrated that P1P treatment during porcine IVM had beneficial effects on oocyte meiotic maturation and developmental competence. During IVM, P1P influenced oocyte maturation as evidenced by the dose-dependent increase in metaphase II rate in the presence of EGF. Evidence also points to reduced ROS levels and alterations in signaling pathways contributing to oocyte development as a result of P1P treatment. Pre-implantation development of oocytes influenced by P1P signaling during IVM also further corroborated evidence of improved developmental competence by P1P treatment.

S1P receptors are a family of G-protein coupled receptors (S1PR1-S1PR5) widely found in multiple organs/systems and during development. S1PR1, S1PR2, and S1PR3 are extensively distributed in various tissues, while S1PR4 and S1PR5 are restrictedly expressed in the immune system and nervous system, respectively [46]. S1PR activation by ligand binding signals downstream Ras/ERK, PI3K/Akt, and Rho/Rock pathways [47–49]. P1P is a ligand able to bind to S1PR1 and S1PR4 with high affinity [11, 12]. In the present study, *S1PR2* and *S1PR3* transcripts were detected in both cumulus cells and oocytes, whereas *S1PR1* transcript was identified only in cumulus cells. S1PR1 and S1PR3 signal downstream pathways via PI3K/Akt exerting anti-apoptotic effects and also via ERKs commonly associated with nuclear maturation and cumulus expansion [33, 50]. Albeit through lower affinity, P1P treatment may also lead to S1PR2 activation previously shown to activate p38 MAPKs and Rho-associated protein kinases (ROCKs) involved in meiotic resumption and spindle positioning, respectively [51–54]. The expression profile of S1PRs on porcine oocytes has suggested that P1P may influence oocyte meiotic progression and developmental competence through interactions with S1PR, possibly initiating similar signaling pathways as seen with S1P supplementation.

To assess the effects of P1P on porcine oocyte meiotic progression, we supplemented P1P during IVM and demonstrated that P1P concentrations between 100 nM and 1000 nM significantly improved meiotic resumption. Our results in P1P supplementation were consistent to a previous study reporting significant differences in meiotic resumption by S1P supplementation during porcine IVM [31]. In contrast, in bovine and murine studies, S1P did not affect meiotic maturation, possibly in part due to the presence of S1P-containing serum or serum-derived products in maturation media providing redundant signaling pathways for the studies to accurately assess the effects of S1P independently. For this reason, we supplemented PVA instead of FF or serum during IVM to minimize the interference from endogenous S1P in FF or serum [25, 55]. Together, our findings suggest that P1P influences meiotic maturation during porcine IVM perhaps through interactions with S1PRs.

In most porcine IVM media, including IVM media used in this study, EGF is supplemented to improve meiotic maturation by influencing cumulus expansion. A number of the previous studies have shown that S1P and P1P have a synergistic effect with growth factors including EGF [34, 35, 56]. In this study, we demonstrated that meiotic maturation was affected by P1P only in the presence of EGF and no significant differences were observed by P1P treatment in the absence of EGF. These results are in agreement with previous studies, one of which also reports a higher level of the p-Akt and p-ERK by combined treatment of EGF and P1P compared to groups with either factor alone [35]. In agreement, western-blot analyses in our study showed increased p-Akt and p-ERK in the presence of P1P and EGF in cumulus cells compared to EGF alone. Interestingly, in an alternate study, treatment of S1P with EGF during porcine IVM have not shown significant differences in meiotic maturation rate while significant effects were observed only in the absence of EGF [31]. It is possible that significant differences observed in the current study may be due to the use of PVA as a serum and FF substitute, allowing for a more independent assessment of the synergistic relationship between P1P signaling and EGF.

In oocytes and embryos, the cytoplasmic GSH level is indicative of cytoplasmic maturation and developmental potential [57]. Often negatively correlated to low GSH levels, high ROS levels have been associated with DNA damage, lipid peroxidation, the expression of particular embryonic genes and meiotic arrest in the oocytes [58, 59]. We have demonstrated that P1P supplementation during IVM lead to a non-significant variation in GSH levels, yet led to a significant decrease in ROS levels. In various biological systems, enzymatic (SOD, CAT, and APX) and non-enzymatic antioxidants (GSH, Vitamin C and beta-carotene) play important roles in controlling ROS to protect oocytes from OS. Since ROS levels were reduced without changes in GSH levels, it may be possible that P1P is reducing ROS levels via regulation of enzymatic antioxidants. In accord, analyses of mRNA expression have confirmed increased expression of enzymatic oxidants such as *SOD3* and *Cat* in oocytes following P1P supplementation. SOD converts superoxide radicals to hydrogen peroxide, and Cat degrades hydrogen peroxide into water and oxygen. Like observed in this study with P1P, S1P also mediates similar effects in cellular antioxidant defense capacity by modulating the activity of enzymatic oxidants (SOD and CAT) and suppressing ROS generation via S1PR2 [60, 61]. Together, these combined observations indicate that P1P improves the quality of IVM oocytes by reducing oocyte intracellular ROS levels possibly via similar mechanisms as S1P. However, it is not entirely clear whether the observed anti-oxidative effects are through direct actions on oocyte by P1P or via cumulus cells known to protect oocytes from OS and express S1PRs [62–64].

To investigate the molecular pathways altered by P1P supplementation in the observed meiotic maturation improvement, we analyzed the expression of select genes associated with extracellular matrix (ECM), enzymatic antioxidant, and apoptosis in cumulus cells and oocytes. ECM modification during cumulus expansion is an essential step during oocyte maturation, influencing developmental competence acquisition downstream of LH surge-induced EGF signaling [65–67]. In post-IVM cumulus cells, *EGF* and *Has2* transcript levels in the P1P treatment group were significantly higher than control groups, while *EGFR and Cox2* transcript levels did not show a significant difference. Several studies have proposed that S1P transactivation or S1PR activation induce EGFR expression and the release of EGF, bearing similar results as in the current study [36, 37, 68]. Downstream of EGF signaling, Cox2 and Has2 expression levels are well-established indicators of the degrees of cumulus expansion and oocyte maturation [69–71]. Specifically, *Has2* regulates the synthesis of hyaluronan, a major structural macromolecule in the cumulus ECM serving as an antioxidant for scavenging ROS [72]. It is possible that P1P-induced increase in EGF expression has led to the increased expression of *Has2*, likely contributing to the decreased ROS and increased developmental competence observed in this study. It may also be possible that the increased p-ERK1/2 and p-Akt observed may be directly responsible for the increased *Has2* expression since ERK 1/2 and PI3K/Akt signaling pathways have been shown to be important for cumulus expansion for oocyte maturation [73–75]. However, despite studies reporting redundant pathways shared between S1P and P1P, our studies using P1P did not show significant changes to *EGFR* and *Cox2* transcript levels, suggesting a possible difference in the molecular pathways activated between S1P and P1P, though further investigations are required [36, 76]. Together, we suggest that P1P exerts positive effects on porcine oocyte maturation and quality in part via increased EGF signaling regulating ERK 1/2 and Akt signaling to influence ECM formation.

Apoptosis plays a crucial role in development, and especially during IVM, increasing numbers of apoptotic cells with DNA fragmentation are observed [77]. In this study, we checked the expression of anti-apoptotic genes (Bcl-2 and Mcl1) and pro-apoptotic genes (Bax and Cas3). In post-IVM oocytes, the expression of anti-apoptotic gene *Bcl-2* was increased and pro-apoptotic gene *Bax* was decreased by P1P exposure. In agreement, previous studies report that exogenous S1P up-regulates the expression of Bcl-2 and Mcl1, while down-regulating the expression of Bax [78–80]. However, in cumulus cells, Bax expression was increased in response to P1P supplementation in our study. Regardless, the *Bcl-2* to *Bax* ratio indicating the balance towards cell survival was improved in cumulus cells as a disproportionately higher increase in *Bcl-2* expression level was observed [41–43]. Increased expression of bcl-2 by P1P may be possibly mediated via increased Akt and ERK phosphorylation shown in our study as also supported by past studies in other systems [81, 82]. Increased Akt signaling is often associated with increased survival by the repression of JNK activation [22]. Often induced by oxidative stress, JNK induces apoptosis in cumulus cells, negatively influencing progesterone production and oocyte developmental competence [83]. We observed that the phosphorylated JNK was decreased in P1P treated cumulus cells, likely meaning partial inactivation of the JNK pathway that positively influenced survival in our study. Programmed cell death or apoptosis is often associated with caspase signaling networks. Several papers demonstrate that S1P inhibits activation of caspase [84, 85]. On the contrary, P1P has not been shown to involve changes in caspase activity [32]. In agreement, Cas3 expression level did not significantly decrease even though upstream pro-apoptotic Bcl2 expression level increased. Also, Mcl1 mRNA levels, which is previously reported to be influenced by caspase-dependent mechanism [86], was not affected in our study. Although caspase mRNA level can only reflect the amount of procaspases and not the level of biological activeness, the quantitative amount of maternal transcripts of caspase-3 has also been shown to play important roles in the caspase cascade of apoptosis [87]. Thus, we suggest that unlike S1P, P1P exhibits an anti-apoptotic effect without altering the expression level of Cas3. Taken together, exogenous P1P supplementation modulates gene expression and signaling pathways involved in the balance between cell death and survival, favoring the latter.

To determine whether the oocytes influenced by P1P also increased in developmental competence, PA and IVF experiments were conducted using oocytes matured in different P1P concentration during IVM. From our experiment comparing cleavage (CL) rate, blastocyst (BL) rate, total cell number, and proportion of apoptotic cells in BL, we deduced that oocytes matured in 1 μM P1P held the highest developmental potential in both quantitative and qualitative aspects. S1P treatment during IVM increased meiotic maturation and BL rate, but unlike P1P, did not affect the total cell number of blastocyst [31, 32]. This may again be linked to the use of serum or FF during IVM in previous observations. Besides P1P’s mechanism in reducing oxidative stress, P1P also upregulated expression of Oct4, a key factor downstream of ERK signaling present in MII oocytes regulating the transition from a gametic to an embryonic development [88, 89].

In summary, the present studies suggest that P1P supplementation in the presence of EGF during porcine IVM improves meiotic resumption and oocyte quality possibly as a result of modulating the activities of several signaling molecules regulating the expression of genes implicated in cumulus expansion, anti-oxidant, developmental competence, and anti-apoptosis. As the positive outcomes suggest, P1P is suitable as a candidate molecule to exploit in the field of reproductive biotechnology.

## Acknowledgments

This work was supported by “Korea Institute of Planning and Evaluation for Technology in Food, Agriculture, Forestry and Fisheries (IPET) through Agri-Bio industry Technology Development Program, funded by Ministry of Agriculture, Food and Rural Affairs (MAFRA) (grant number: 318016-5)” and by “Business for Cooperative R&D between Industry, Academy, and Research Institute funded Korea Small and Medium Business Administration in 2017 (Grants No. 2017020681010101).”

## Author Contributions

Conception and design of study: KMP YWJ KCH WSH MJC SHH EBJ. Acquisition of data: KMP YMY JEJ. Analysis and/or interpretation of data: KMP YMY KCH JWW. Writing: KMP JWW. Critical revision: KMP YWJ JWW

---

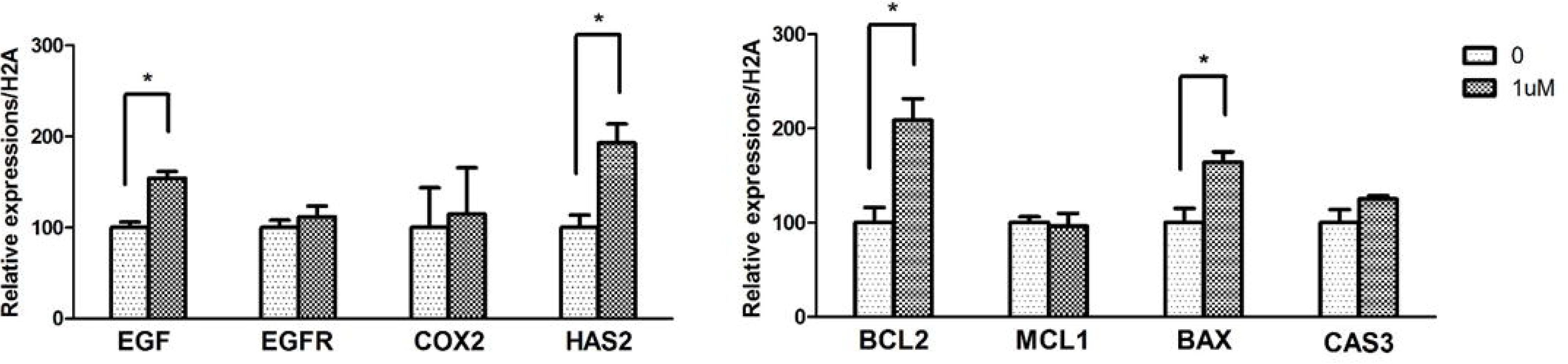

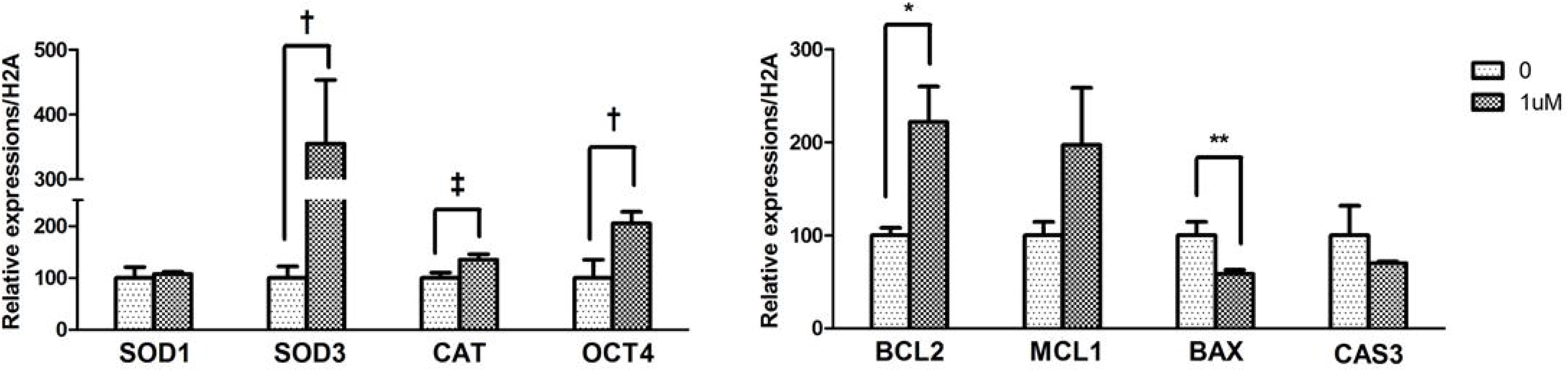

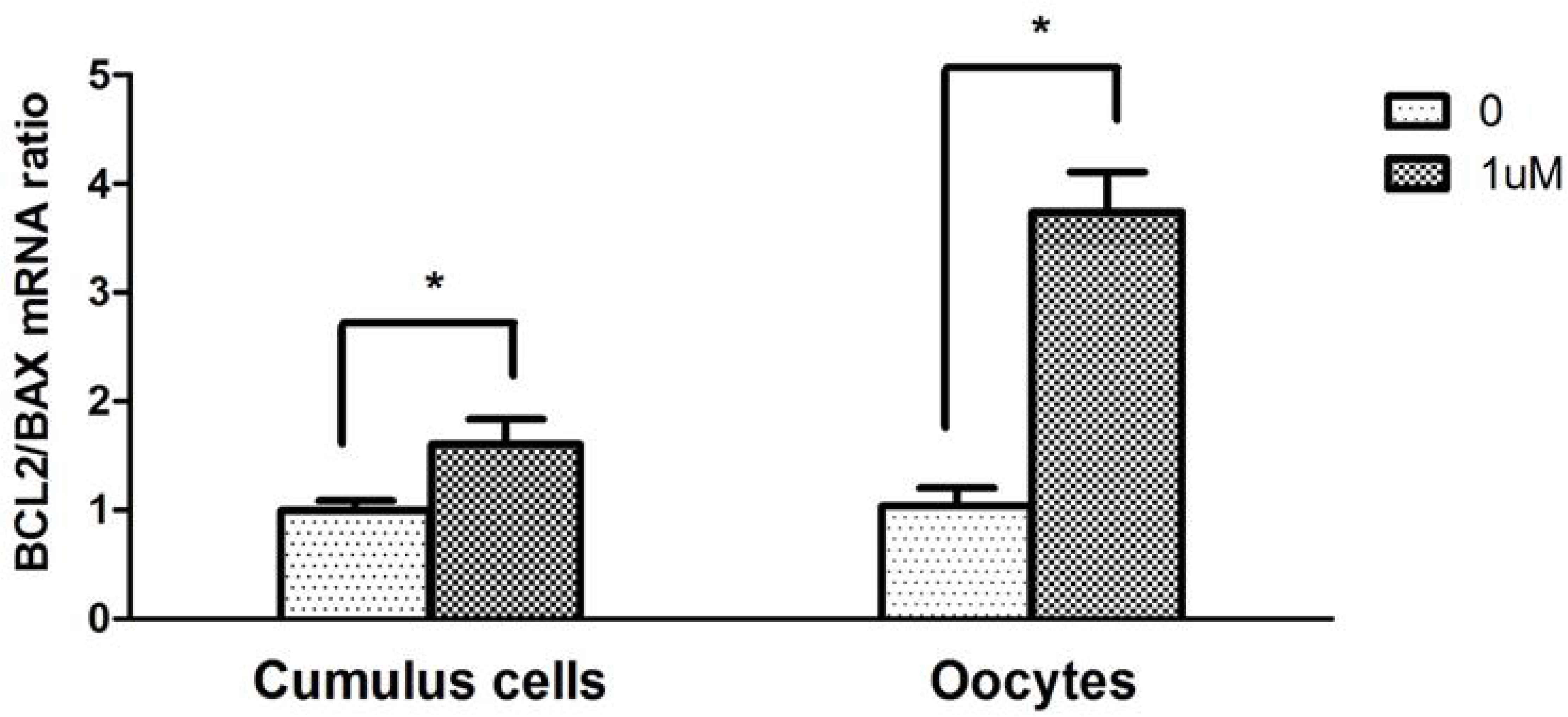

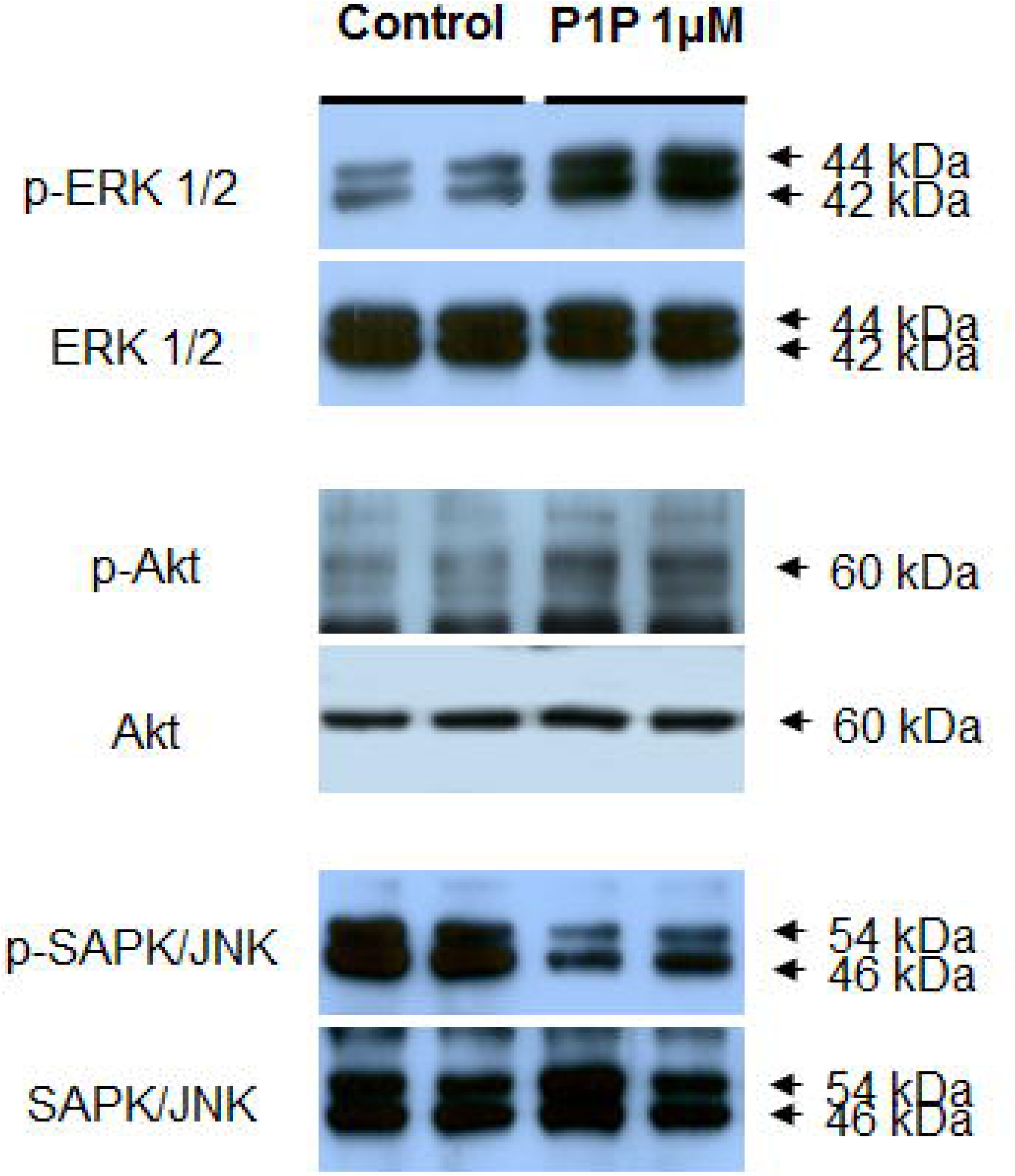

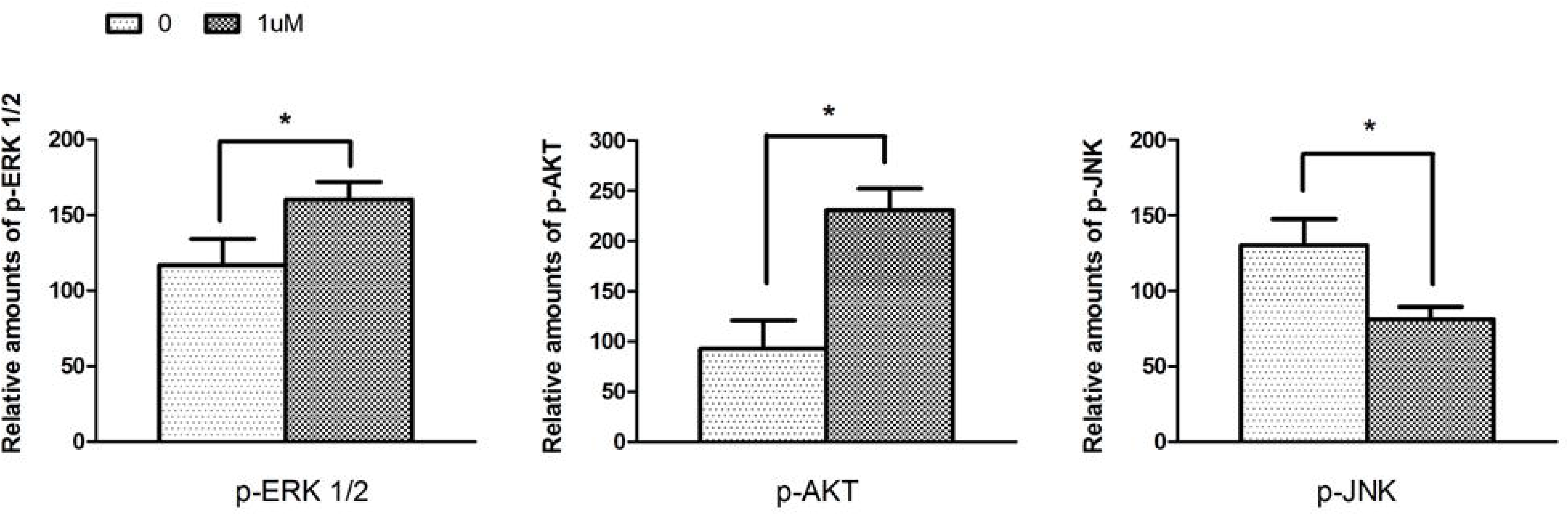

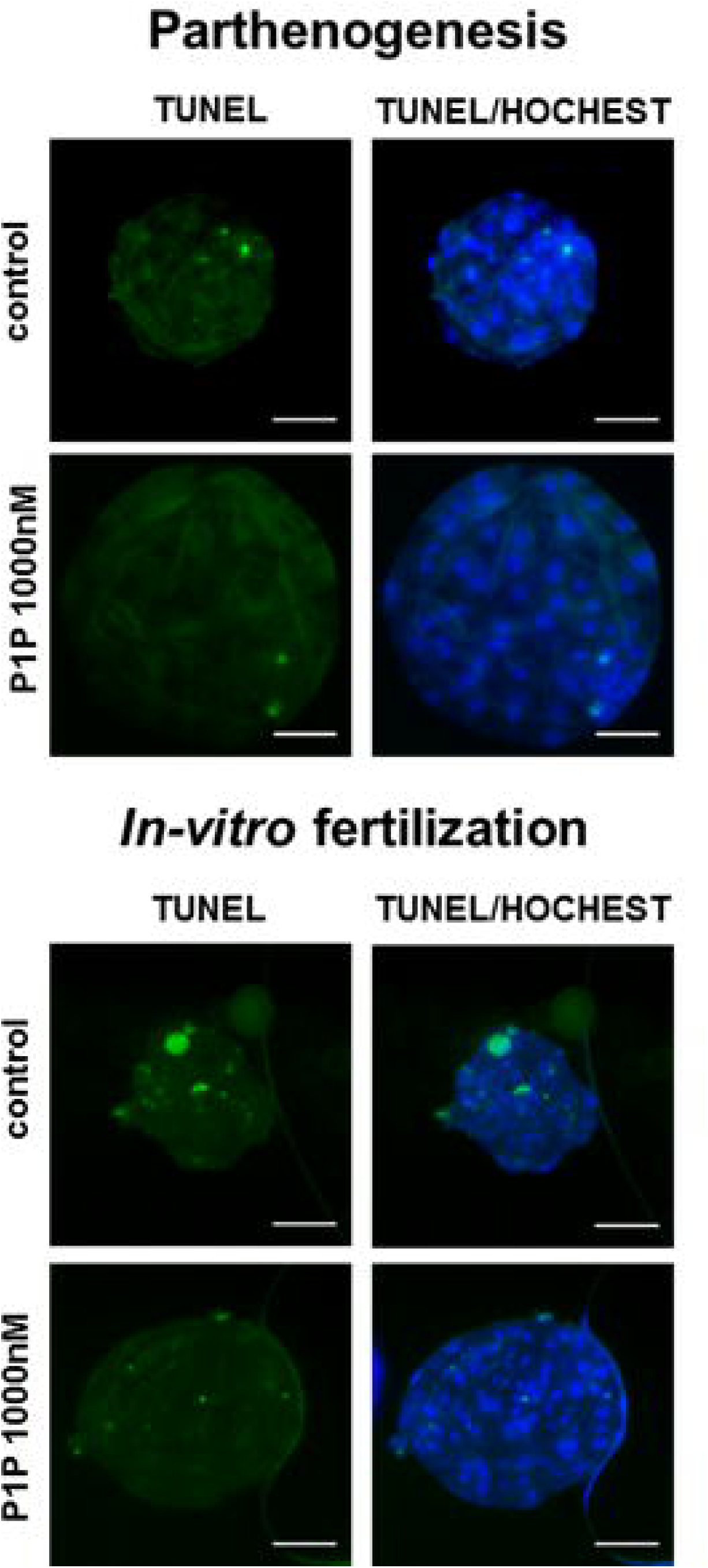

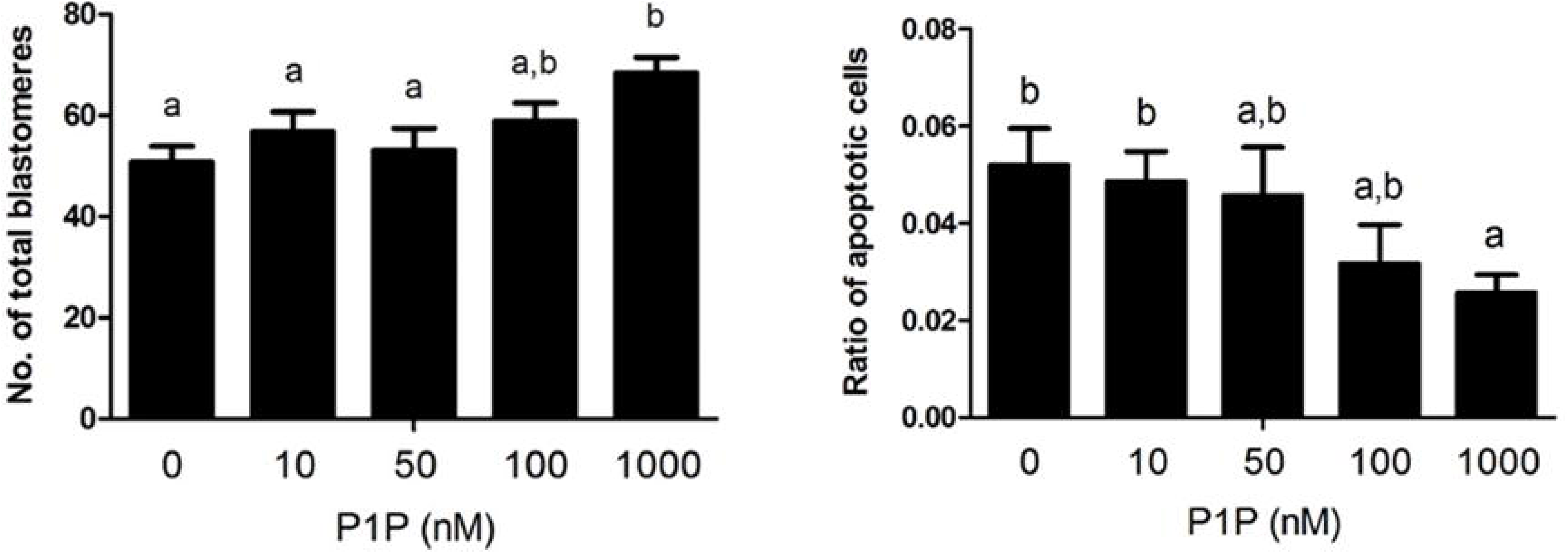

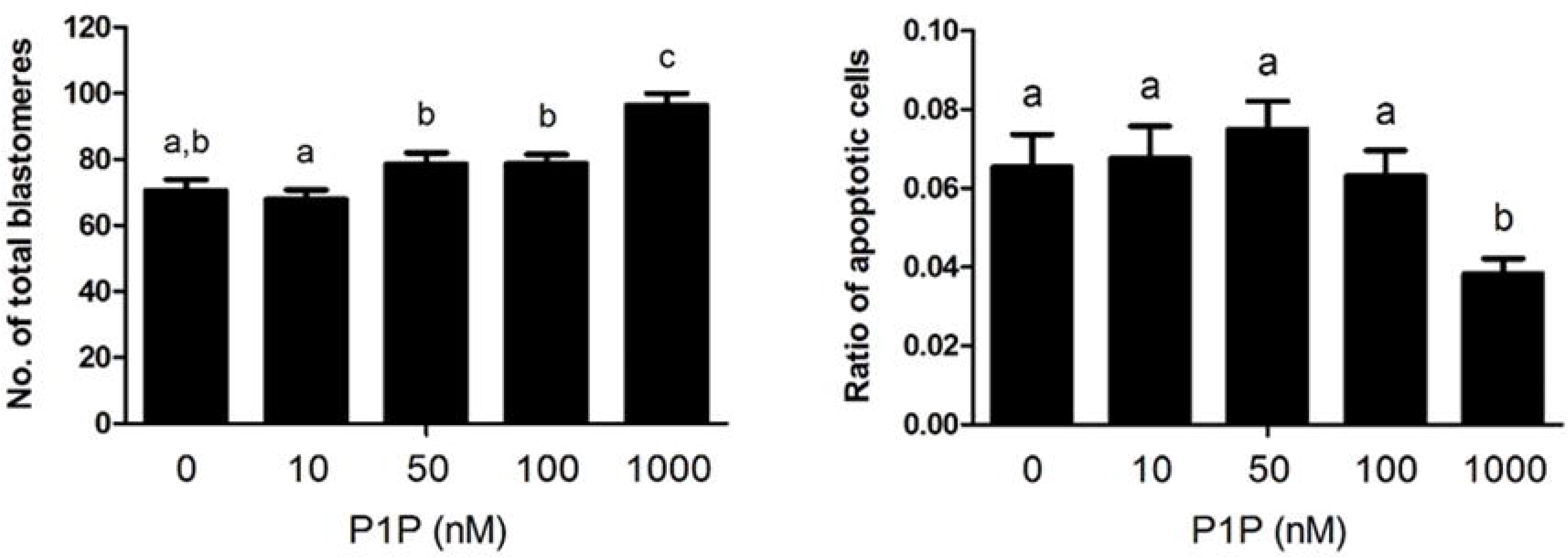

