## supplemental table 1 for "Sphingosine-1-Phosphate (S1P) analogue Phyto-sphingosine-1-Phosphate (P1P) improves the *in vitro* maturation efficiency of porcine oocytes via regulation of oxidative stress and apoptosis"

**S1 Table.** Concentration-dependent effects of P1P treatments on nuclear maturation in porcine oocytes

| P1P  concentration  (μM) | No. of oocytes  cultured for IVM(4)* | No. (%) of oocytes at the stage of | | | | |
| --- | --- | --- | --- | --- | --- | --- |
|  |  | GV | GVBD | MI | AI/TI | MII |
| 0 | 234 | 17(7.3±2.0)^b^ | 0(0.0) | 20(8.5±1.6) | 2(0.8±0.8)^a^ | 195(83.3±1.2)^a^ |
| 1 | 238 | 7(2.9±1.3)^a,b^ | 0(0.0) | 14(5.9±1.5) | 0(0±0.0)^a^ | 217(91.2±2.7)^b^ |
| 5 | 239 | 8(3.3±2.0)^a^ | 0(0.0) | 24(10.1±1.6) | 17(7.1±1.0)^b^ | 190(79.5±1.7)^a^ |

The data are mean±SEM

*Replication number

^a,b^Values with different superscripts within a column differ significantly(p<0.05)

Abbreviation: AI/TI, anaphase I/telophase I; GV, germinal vesicle; GVBD, germinal vesicle breakdown; MI, metaphaseI; M II, metaphase II
